# Testing the causal effects between subjective wellbeing and physical health using Mendelian randomisation

**DOI:** 10.1101/304741

**Authors:** Robyn E Wootton, Rebecca B Lawn, Louise A C Millard, Neil M Davies, Amy E Taylor, Marcus R Munafò, Nicholas J Timpson, Oliver S P Davis, George Davey Smith, Claire M A Haworth

**Affiliations:** School of Experimental Psychology, University of Bristol, Bristol, BS8 1TU; Department of Population Health Sciences, Bristol Medical School, University of Bristol, Bristol, BS8 2PR; MRC Integrative Epidemiology Unit, University of Bristol, Bristol, BS8 2PR; NIHR Biomedical Research Centre at the University Hospitals Bristol NHS Foundation Trust and the University of Bristol, (BS8 2BN).; UK Centre for Tobacco and Alcohol Studies, University of Bristol, Bristol, BS8 1TU; Intelligent Systems Laboratory, Department of Computer Science, University of Bristol, Bristol, BS8 1UB

## Abstract

**Objectives:** To investigate whether the association between subjective wellbeing (subjective happiness and life satisfaction) and physical health is causal.

**Design:** We conducted two-sample bidirectional Mendelian randomisation between subjective wellbeing and six measures of physical health: coronary artery disease, myocardial infarction, total cholesterol, HDL cholesterol, LDL cholesterol and body mass index (BMI).

**Participants:** We used summary data from four large genome-wide association study consortia: CARDIoGRAMplusC4D for coronary artery disease and myocardial infarction; the Global Lipids Genetics Consortium for cholesterol measures; the Genetic Investigation of Anthropometric Traits consortium for BMI; and the Social Science Genetics Association Consortium for subjective wellbeing. A replication analysis was conducted using 337,112 individuals from the UK Biobank (54% female, mean age =56.87, SD=8.00 years at recruitment).

**Main outcome measures:** Coronary artery disease, myocardial infarction, total cholesterol, HDL cholesterol, LDL cholesterol, BMI and subjective wellbeing.

**Results:** There was evidence of a causal effect of BMI on subjective wellbeing such that each 1 kg/m^2^ increase in BMI caused a 0.045 (95%CI 0.006 to 0.084, *p*=0.023) SD reduction in subjective wellbeing. Replication analyses provided strong evidence of an effect of BMI on satisfaction with health (β=0.034 (95% CI: −0.042 to −0.026) unit decrease in health satisfaction per SD increase in BMI, *p*<2^-16^). There was no clear evidence of a causal effect between subjective wellbeing and the other physical health measures in either direction.

**Conclusions:** Our results suggest that a higher BMI lowers subjective wellbeing. Our replication analysis confirmed this finding, suggesting the effect in middle-age is driven by satisfaction with health. BMI is a modifiable determinant and therefore, our study provides further motivation to tackle the obesity epidemic because of the knock-on effects of higher BMI on subjective wellbeing.

## Introduction

Subjective wellbeing is most commonly defined as a combination of life satisfaction and positive affect in the absence of negative affect [1]. Observational evidence suggests an association between higher subjective wellbeing and improved physical health or longevity [2–4], including cardiovascular outcomes [5], cholesterol levels [6] and extremes of body mass index (BMI) [7].

Depression has been shown to have the opposite association with physical health, increasing risk of coronary artery disease (CAD) especially the chance of a heart attack [8], altering serum cholesterol [9] and a U-shaped relationship with BMI [10]. A Mendelian randomisation (MR) study of BMI on multiple mental health outcomes found a consistent effect of higher BMI on increased likelihood of depression, although the effect sizes were small [11]. This causal effect was replicated in the follow-up analysis of the most recent genome-wide association study (GWAS) of depression [12] and was replicated using a continuous measure of depressive symptoms [13], with suggestive evidence of a causal link between BMI and subjective wellbeing. However, this study did not examine other health behaviours and did not adjust for sample overlap, so results could be biased towards the observational effect [14].

Twin analyses indicate partly distinct genetic (and environmental) aetiologies for depression and subjective wellbeing [15], suggesting that separate analyses of the relationship between subjective wellbeing and depression on health outcomes may be appropriate. Observational research suggests that the association between subjective wellbeing and physical health goes beyond the absence of negative affect states, reduced arousal or confounding from socio-economic position [16] and subjective wellbeing is more predictive of health outcomes than negative feelings [17]. Therefore, subjective wellbeing might be a better target for improving physical health outcomes than depression. From a public health perspective, it is important to understand whether increasing subjective wellbeing can increase health in later life, given that wellbeing interventions can be cost-effective to administer [18].

Studies suggesting a link between subjective wellbeing and physical health are mostly observational. Due to reverse causation and residual confounding, it is hard to make causal inferences from observational evidence [19]. Mendelian randomisation (MR) uses genetic variants as instrumental variables for the exposure of interest. Mendelian randomisation exploits Mendel’s laws of segregation and independent assortment: the inheritance of the alleles will be largely independent of genetic variants affecting other traits and of conventional disease risk factors. Associations are not affected by reverse causation, because genotype is unchanged over the lifetime [19,20]. In an instrumental variable analysis, the genetic variant (Z) acts as the instrument which is related to differences in the exposure (X). If the exposure causes the outcome (Y) then genetic variants which affect the exposure should be associated with the outcome (see Figure 1) [19]. For example, genetic variants (Z) shown to predispose individuals to have a higher BMI (X) are associated with lower income, suggesting increases in BMI reduce income (Y) [21].

**Figure 1.**
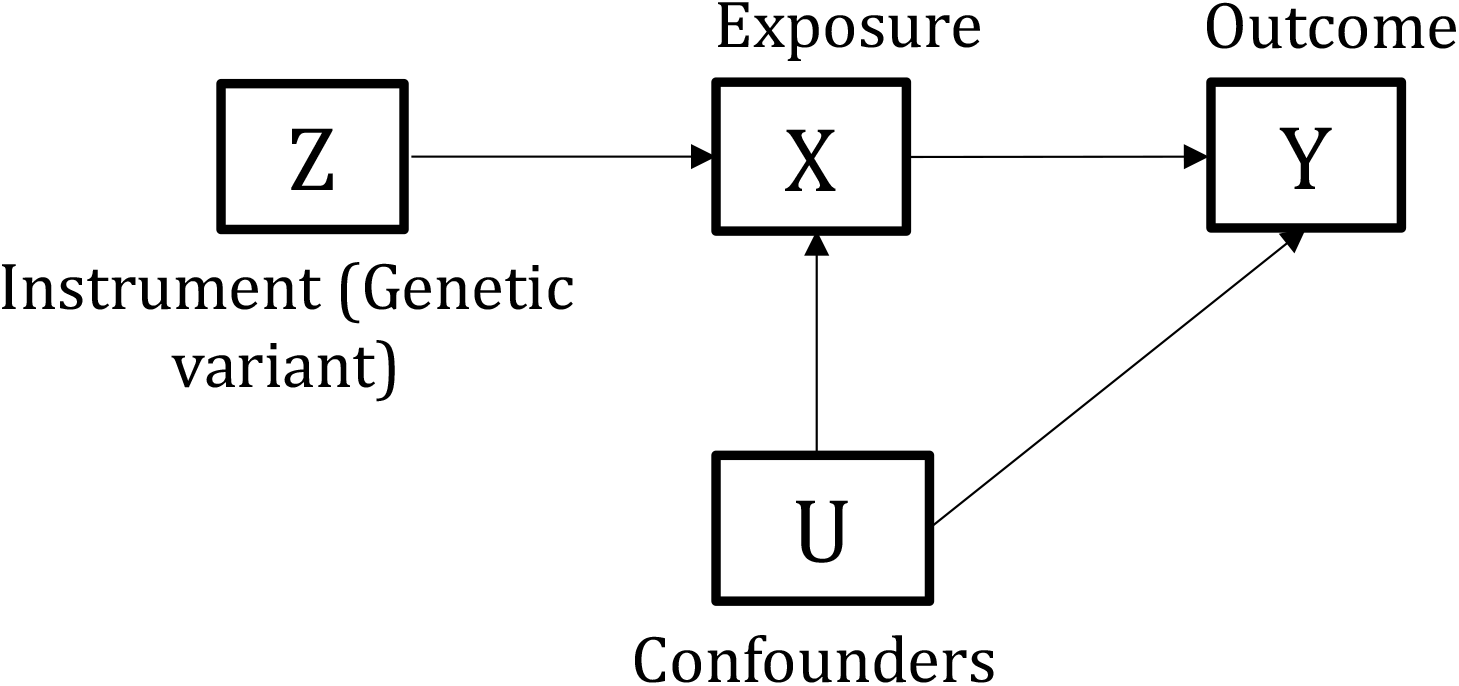
A Directed Acyclic Graph (DAG) representing the basic Mendelian Randomisation instrumental variable analysis.

The two-sample MR method uses the summary statistics from two separate Genome-wide Association Studies (GWAS) in one analysis [22]. The GWAS of the exposure must have identified single nucleotide polymorphisms (SNPs) robustly associated with the exposure. These SNPs can then be looked up in the GWAS summary statistics of the outcome. For power, multiple genetic variants, rather than single genetic variants, are often used in two-sample MR. However, this increases the likelihood of pleiotropic effects biasing the result, so it is particularly important to perform sensitivity analyses that are more robust to pleiotropy when using multiple instruments [14].

Pleiotropy occurs when a genetic variant affects multiple phenotypes. There are two classes of pleiotropy of relevance to MR: horizontal and vertical [22]. Vertical pleiotropy occurs when one genetic variant has multiple effects all on one causal pathway. Vertical pleiotropy does not violate the assumptions of MR as long as the phenotype most proximal to the genetic variation is correctly identified [22]. Horizontal pleiotropy occurs when a genetic variant affects the outcome via pathways aside from through the exposure of interest [22] and can bias MR estimates [19].

Mendelian randomisation makes several assumptions that must be checked to ensure the validity of the causal conclusions. First, the genetic instrument (Z) must be associated with the exposure (X) in the sample. Second, the genetic instrument must not associate with the confounders (U). Finally, Z has no effects on the outcome apart from through the exposure X [19].

In this study, we investigated the association between subjective wellbeing and the physical health traits of BMI, coronary artery disease (CAD), HDL, LDL and total cholesterol and Myocardial Infarction (MI) using MR. We conducted two-sample bidirectional MR analyses to establish whether subjective wellbeing affects physical health traits, or vice-versa. We extended previous research by conducting follow up analysis in an independent sample to avoid sample overlap [14] and examining the causal relationship between subjective wellbeing and a range of physical health conditions.

## Method

### Data Sources

Details of the contributing GWAS consortia are given in Table 1. They were selected for traits relating to cardiovascular health or obesity, having the largest sample size and consisting of the most similar populations.

**Table 1.** Description of GWAS consortia used for each phenotype

| Variable | First Author | Year | Consortium | Sample size | Population | Gender |
| --- | --- | --- | --- | --- | --- | --- |
| **Subjective Wellbeing** | Okbay [33] | 2016 | SSGAC | Exposure = 298420<br>Outcome = 197174 | European | Mixed |
| **Coronary Artery Disease** | Nikpay [36] | 2015 | CARDIoGRAMplusC4D | Cases = 60801<br>Controls = 123504 | Mixed | Mixed |
| **Total Cholesterol** | Willer [37] | 2013 | GLGC | 92260 | Mixed | Mixed |
| **HDL Cholesterol** | Willer [37] | 2013 | GLGC | 92860 | Mixed | Mixed |
| **LDL Cholesterol** | Willer [37] | 2013 | GLGC | 83198 | Mixed | Mixed |
| **Myocardial Infarction** | Nikpay [36] | 2015 | CARDIoGRAMplusC4D | Cases = 43676<br>Controls = 128199 | Mixed | Mixed |
| **BMI** | Locke [34] | 2015 | GIANT | 339224 | Mixed | Mixed |

*Note: SSGAC = Social Science Genetics Association Consortium, GLGC = Global Lipids Genetics Consortium, GIANT = The Genetic Investigation of Anthropometric Traits consortium. Sample sizes were different when wellbeing was the exposure or outcome due to 23andMe data only being included for the top 10,000 SNPs. Therefore, data including 23andMe was used for the exposure, but the data without 23andMe was used for the outcome.*

### Statistical Analyses

We applied four different two-sample MR methods, which make different assumptions about horizontal pleiotropy. Therefore, a consistent effect across the four methods is less likely to be a false positive [23]. If the genetic variants have horizontally pleiotropic effects but they are independent of the effects of the genetic variants on the exposure, then this is known as balanced pleiotropy. If all of the pleiotropic effects are biasing the estimate in the same direction (directional pleiotropy) this will bias the results (with the exception of the MR Egger method). We used the MR Egger intercept to test for the presence of directional pleiotropy.

#### Instrument identification in MR Base

For all phenotypes, apart from subjective wellbeing, our genetic instruments were composed of genome-wide significant SNPs (*p*<5×10^−8^) from published GWAS studies. Only three genome-wide significant SNPs were available for subjective wellbeing. We tested the strength of these instruments by checking if they predicted happiness in a large independent sample (N= 242,219) from the UK Biobank. There was only evidence that one SNP (rs2075677) was associated with happiness (see supplementary Table S1). Therefore, we used a more liberal *p*-value threshold of *p*<5×10^−5^ as the instrument for subjective wellbeing. SNPs were clumped to ensure independence at LD R^2^ = 0.001 and a distance of 10000kb. If a SNP was unavailable in the outcome GWAS summary statistics, then proxy SNPs were searched for with a minimum LD R^2^ = 0.8. MAF was used to align palindromic SNPs with MAF<0.3. Inverse-variance weighted, MR-Egger, weighted median and weighted mode approaches were compared. Analyses were conducted using MR-Base [24], a package for two-sample MR.

#### Inverse-variance weighted method

The inverse-variance weighted (IVW) meta-analysis uses the individual Wald ratios conducted for each SNP. The Wald ratio is calculated by regressing each genetic variant on the exposure and outcome separately. When effects on the outcome and exposure are plotted, the gradient of the line of best fit taking into account all of the data points (constrained to have an intercept of 0) gives the strength of the association. The slope indicates how much of a unit increase there is in exposure relative to each unit increase in outcome [25]. This method may be biased by horizontal pleiotropy [26].

#### MR-Egger method

The MR-Egger method relaxes the assumptions of MR and allows for directional pleiotropic effects, such that some SNPs could be acting on the outcome through a pathway other than through the exposure. Unlike the IVW method, the intercept is no longer constrained to pass through zero. This allows an adjustment to be made for the presence of directional pleiotropy. The intercept itself is an estimate of the directional pleiotropic effect [26]. MR-Egger has the lowest power of the four methods to detect a causal effect, and requires variation in the SNP effects, and therefore is more effective when more SNPs are used to create the instrument. The MR-Egger method also makes the additional NOME assumption, that there is no measurement error in the SNP-exposure effects [26]. This is evaluated using the I^2^_(GX)_ statistic [27].

#### Weighted median method

Rather than relaxing the assumptions of pleiotropy for all SNPs used (like MR-Egger), the weighted median approach assumes that at least 50% of the total weight of the instrument comes from valid variants. The weighted median approach is more likely to give a valid causal estimate than MR-Egger or IVW because it is more consistent with the true causal effect in the presence of up to 50% invalid variants [28].

#### Weighted mode-based estimation method

The weighted mode-based estimation (MBE) method assumes that the most common causal effect is consistent with the true causal effect [29] Hence, the remaining instruments could be invalid (violate the assumptions of MR) without biasing the estimated causal effect.

### Replication in the UK Biobank

We attempted replication of our two-sample MR results (to overcome potential bias from sample overlap) using MR analysis where participants for the exposure and outcome were from the same sample (UK Biobank), with a weighted genetic score derived using estimates from GWAS. The replication sample and measures are now described below.

### Study sample

UK Biobank is a national health resource with biological measures collected over 10 years (http://www.ukbiobank.ac.uk). A total of 502,647 participants aged 40-69 years were recruited from across the United Kingdom between 2006 and 2010 [30]. After restricting to European ancestry and excluding related individuals, withdrawn consent and sex mismatched individuals, 337,112 participants remained [31]. Of these individuals, the mean age was 56.87 (SD = 8.00) years at recruitment and 54% were female.

A sub-sample (150,000) participants were genotyped first, and this sample was selected on smoking status to include more current smokers than would be representative of the UK population [32]. These 150,000 genotyped individuals also contributed to the SSGAC GWAS of subjective wellbeing [33]. The remaining UK Biobank participants have now been genotyped. To avoid any possible biases associated with smoking, we used the full Biobank sample in the replication analysis presented here (N = 337,112). However, because of partial sample overlap with the SSGAC GWAS, we repeated the same replication analysis including only individuals who did not contribute to the SSGAC GWAS (N = 242,219).

### BMI allele score

To conduct the replication of the link between BMI and wellbeing we constructed a polygenic score for BMI in UK Biobank. This polygenic score was constructed by extracting the 97 variants found to be associated at genome-wide significance with BMI in the most recent GWAS [34]. Allele scores for each SNP were calculated as a sum of the number of increasing alleles weighted by the effect sizes from the GWAS summary statistics. Therefore, higher polygenic score indicates an increased risk of higher BMI. Of the 97 SNPs, rs2033529 was unavailable in UK Biobank (see Supplementary Table S2 for full SNP list). The allele score was standardised to mean zero and standard deviation one.

### Observed BMI

Body Mass Index was calculated (weight in kg/(height in m)^2^) from measurements of height and weight taken during the initial assessment centre visit.

### Outcomes

We used phenotypic measurements collected at initial assessment (2006-2010). The measures were

*Subjective Happiness* - assessed using a single item questionnaire measure. Responses to the question ‘In general how happy are you?’ were collected on a 6-point likert scale ranging from *Extremely unhapppy* to *Extremely happy*. Individuals responding *Do not know* or *Prefer not to answer* were coded as missing.

*Life Satisfaction* - assessed using five single item measures relating to domains of life satisfaction. Domains were: family and relationships, work/job, health, finances and friendships. For example, ‘In general how satisfied are you with your family relationships?’ Responses were collected on a 6-point Likert scale ranging from *Extremely unhapppy* to *Extremely happy*. Individuals could also respond *Do not know* or *Prefer not to answer* (and additionally *I am not employed* for the work/job domain), which were coded as missing.

*Baseline Demographic Measures* - collected at initial assessment, including: sex, age and socio-economic position (SEP). SEP was measured using the Townsend deprivation index (Townsend, 1987) based upon their location in the UK (calculated from output area) and information from the last national census.

### Statistical Analysis

Mendelian randomisation was conducted through instrumental variable regressions run in R [35] to attempt to replicate the effect of BMI on subjective wellbeing using the polygenic score for BMI as the instrument.

## Results

### Two-sample MR: Subjective Wellbeing Predicting Physical Health Outcomes

The genetic instrument for the exposure, subjective wellbeing, was 84 SNPs with *p*<5×10^−5^ (independent at R^2^ = 0.001 and a distance of 10000kb) from the GWAS by the SSGAC [33]. The regression dilution I^2^_(GX)_ estimate was less than 90% for the subjective wellbeing instrument (see Supplementary Table S3 for further information), therefore simulation extrapolation (SIMEX) correction was applied in MR Egger analysis [27]. There was no clear evidence to suggest a causal effect of subjective wellbeing on any of the health outcomes (see Figure 2).

**Figure 2.**
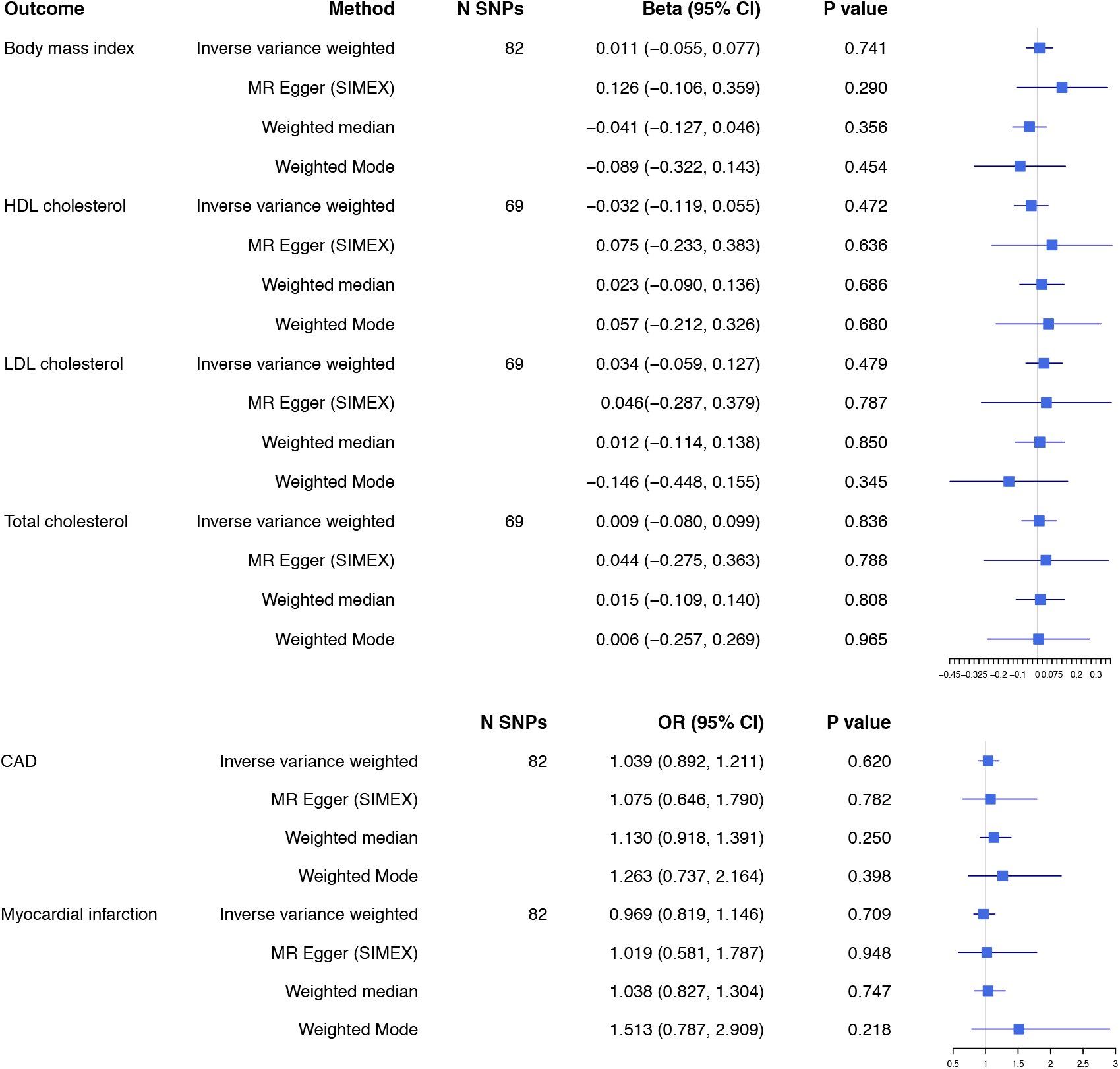
Two-sample MR analysis: the effect of subjective wellbeing on physical health outcomes using SNPs significant at *p*<5×10^−5^. One unit increase of subjective wellbeing is equivalent to one standard deviation increase of the subjective wellbeing composite continuous scale. N SNP refers to the number of the 84 SNPs associated with wellbeing which were available in the outcome summary statistics. SNPs might be unavailable in the outcome due to imputation platform or not passing QC procedures. A more stringent analysis using only genome-wide significant SNPs as the instrument produced a similar pattern of results (see Supplementary Figure S1).

### Two-sample MR: Physical Health Predicting Subjective Wellbeing

In this analysis, we investigated whether subjective wellbeing was causally affected by physical health. Genome-wide significant SNPs for each physical health measure were used as genetic instruments. The number of SNPs this gave for each analysis is given in Figure 3. The regression dilution I^2^_(GX)_ estimates for all exposures were greater than 90% (see Supplementary Table S3 and Supplementary Note for further information), indicating exposures were suitable for MR Egger analysis [27]. We found evidence that higher BMI caused lower subjective wellbeing (see Figure 3). The direction of effect remained consistent across all four methods. There was no clear evidence of a causal effect of any of the heart health or cholesterol exposures on subjective wellbeing (see Figure 3).

**Figure 3.**
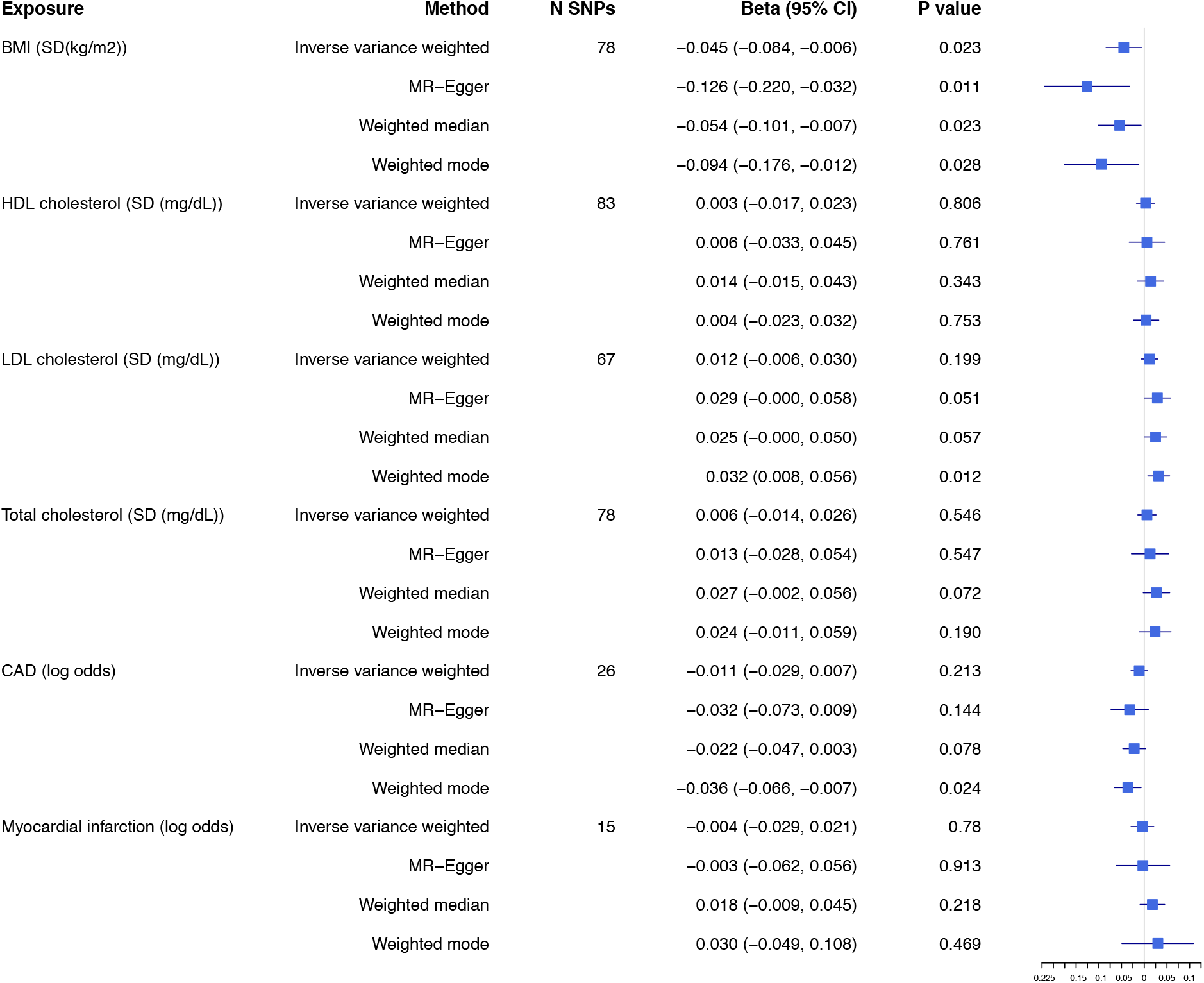
Two-sample MR analysis: the effect of physical health exposures on subjective wellbeing per unit of exposure. N = number of SNPs. For all phenotypes, genome-wide significant SNPs (*p*<5×10^−8^) from the previous GWAS studies were used as the instrument, clumped at LD R^2^ = 0.001 and 10000kb. Genome-wide significant SNPs have odds ratios from 1.04 (1.02, 1.06) - 1.37 (1.31, 1.44) for CAD risk and from 1.03 (1.01, 1.06) - 1.33 (1.27, 1.4) for MI risk [36]. Genome-wide significant SNPs for BMI account for 2.7% of variance in BMI [34], 13.7% of the variance in HDL cholesterol, 14.6% of LDL cholesterol and 15% of total cholesterol [37,38].

Cochran’s Q and I^2^ statistics were calculated to check for the presence of heterogeneity (dispersion of SNP effects), which can indicate pleiotropy. There was little evidence of heterogeneity for the association between BMI and wellbeing (see Supplementary Table S4 for results and further information). The MR-Egger intercept suggested little evidence of directional pleiotropy (see Supplementary Table S5, all p>0.07). The funnel plot of individual SNP effects revealed a symmetrical distribution of SNP effects around the effect estimate suggesting balanced pleiotropy (see Supplementary Figure S2). We also conducted a leave-one-out analysis revealing the SNP with the largest contribution to the effect is rs1421085 located on chromosome 16 in the second intron of the *FTO* (fat mass and obesity associated) gene (see Supplementary Figure S3 and for more information).

### Replication in the UK Biobank

#### Observational association between BMI and subjective wellbeing

Mean BMI in the UK Biobank replication sample was 27.39 (SD=4.75). Means and standard deviations for the subjective wellbeing measures (scored from 1-6 with 6 being high wellbeing) are given in Table 2. Mean subjective wellbeing values show some negative skew but none have skew less than −1. Linear regressions were conducted to test the observational association between BMI and subjective wellbeing in our UK Biobank sample controlling for age, sex and SEP (see Table 2). BMI was negatively associated with all measures of subjective wellbeing apart from job satisfaction and satisfaction with family where there was no clear association and satisfaction with friends where the association was positive.

**Table 2.**
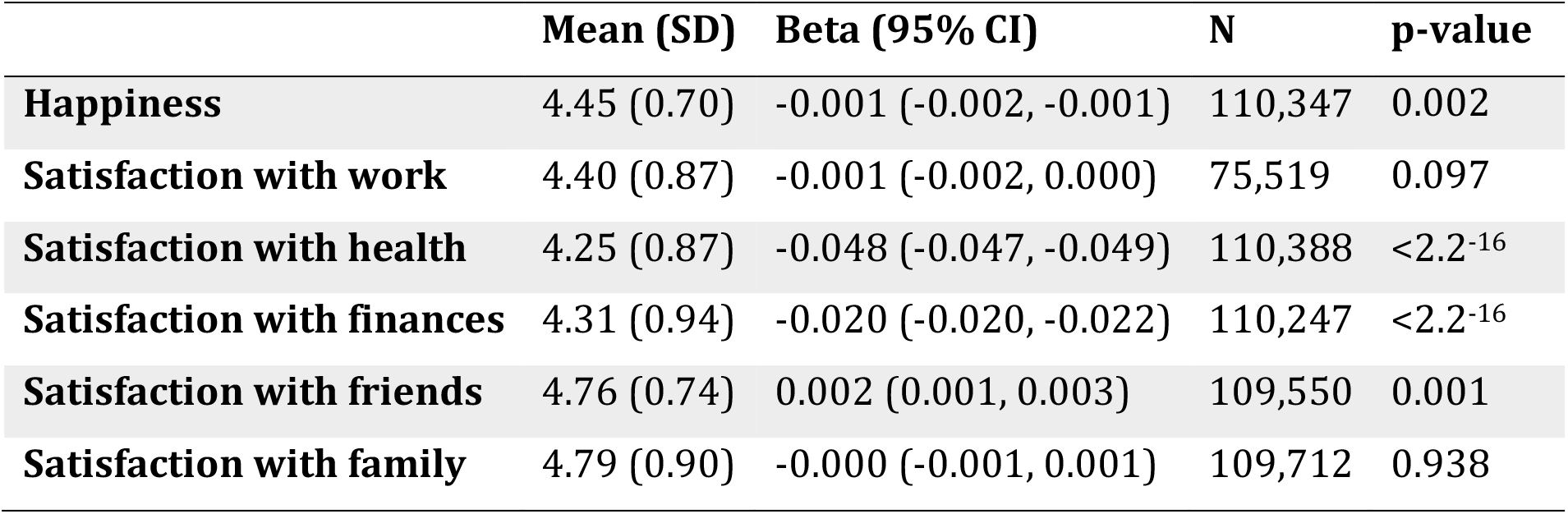
Linear regressions between BMI and subjective wellbeing in the UK Biobank sample

#### Association with baseline confounders

The association of the BMI genetic score and BMI with baseline confounders (age, sex, SEP, education, smoking and alcohol consumption) were compared (see Supplementary Figure S4). There was evidence of an association between our BMI genetic score and SEP, educational attainment, smoking behaviour and alcohol consumption. For educational attainment and SEP, the association was weaker for the genetic score than observed BMI. The association between our BMI genetic score with daily alcohol consumption and smoking could be due to a causal effect of BMI on these outcomes [39–41].

#### Association between polygenic score and BMI

The genetic score was strongly associated with observed BMI (strength of instrument: F(1, 336027) = 6180, R^2^ = 0.018, *p*<2.2^-16^).

#### Replication analysis of BMI (exposure) on subjective wellbeing (outcome)

The results are shown in Figure 4. There was very strong evidence of a causal effect of BMI on satisfaction with health (*β*=-0.034, 95% CI −0.042 to −0.026, *p*<2^-16^). There was little clear evidence of a causal effect of BMI on any of the other measures of subjective wellbeing. There was little evidence that this effect differed in older and younger participants, but the age range in the UK Biobank is narrow with all participants over 40 years old. When individuals were split by median age, the evidence for a causal effect of BMI on satisfaction with health remained strong in both groups (Age≤58 years: −0.040, 95% CI −0.050 to −0.029, *p*=6.4^13^; Age>58 years: −0.028, 95% CI −0.040 to −0.016, *p*=3.7^-6^). The results remained the same in the independent sample with contributors to the subjective wellbeing SSGAC GWAS removed (see Supplementary Table S6).

**Figure 4.**
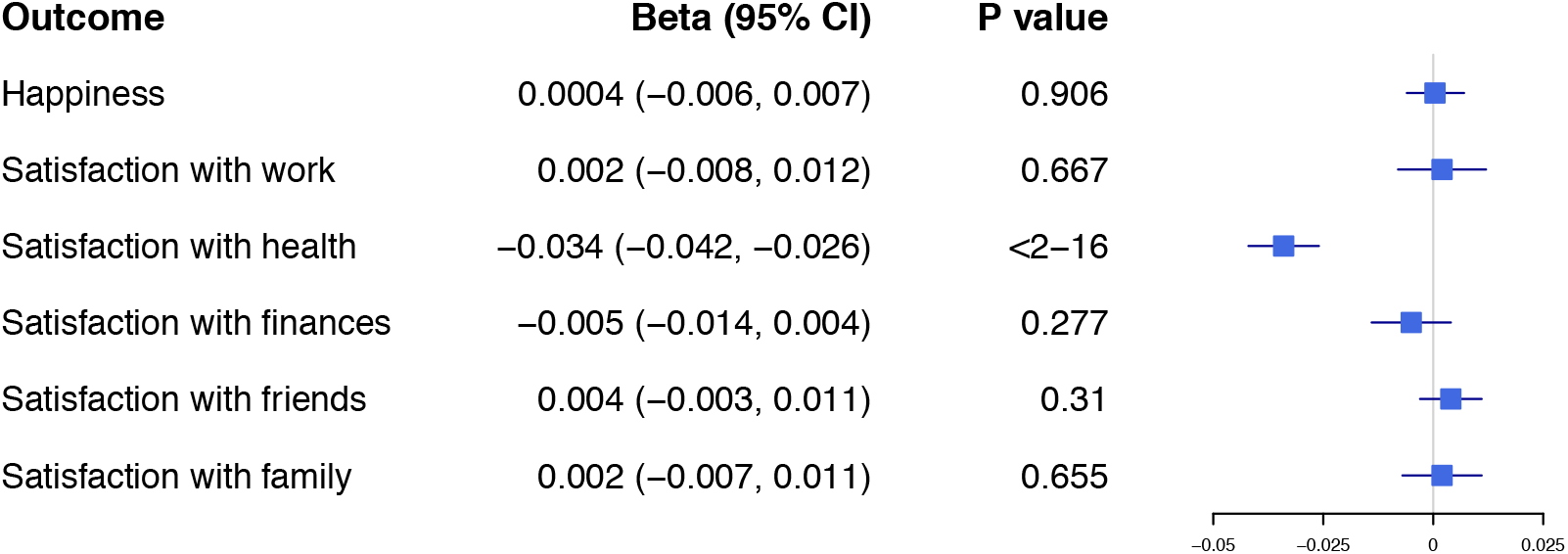
Results of the replication analysis: effect on subjective wellbeing per 1kg/m^2^ of BMI, consistent with previous estimates from automated MR-PheWAS [42].

## Discussion

We found evidence that higher BMI causes lower subjective wellbeing. Sensitivity analyses suggested this was not due to directional pleiotropy and the finding replicated in the UK Biobank. The replication analysis suggested the causal effect of BMI on subjective wellbeing was driven by satisfaction with health, such that higher BMI caused lower health satisfaction. The pathway from BMI to health satisfaction could be biological or social. Biological pathways include BMI as a risk factor for other negative health outcomes such as diabetes, cardiovascular illness and cancers [43] with randomised control trials and MR stengthening evidence of a causal effect [44–46]. Therefore, the effect of BMI on satisfaction with health seen in the current study may reflect accurate perceptions of health. Alternatively, societal influences could cause individuals to associate negative health consequences with a higher BMI and consequently report lower health-satisfaction. Subjective wellbeing and health are in a complex and dynamic system of causal pathways and further work is needed to understand these using mediation analysis [2].

In understanding this causal effect further, another important consideration might be the influence of age. Individuals recruited for the UK Biobank are middle aged or older, with an average age of 57 years. BMI may be an important determinant of health satisfaction in an older generation as the health implications of obesity (heart disease, diabetes, cancer) begin to emerge [47]. In younger age groups, body dissatisfaction, self-esteem and bullying might be more important mediators of the association between BMI and wellbeing [48]. Further investigation of the causal pathways in a younger sample should be explored, especially as some genetic variants for BMI show a developmentally specific pattern of association[49,50].

Observational evidence suggests a non-linear association between mental health and BMI, where extremely high and low BMI both predict higher rates of depression and lower rates of wellbeing [7,10]. The association between very low BMI and depression seen in observational studies could be driven by eating disorders such as anorexia nervosa. The two are commonly comorbid with a 50% lifetime prevalence of major depression in individuals with anorexia [51]. Twin studies have suggested this comorbidity is due to shared genetic influence between anorexia and major depressive disorder [52]. However, the Mendelian randomisation estimators we used assume a linear relationship. Therefore, if individuals with low BMI also have lower subjective wellbeing, this could lead to the effect observed in MR appearing smaller than it truly is. New methods to allow for non-linear associations in MR are being developed [53,54] but are currently too underpowered to apply here.

Two-sample MR analyses revealed no clear evidence of a causal effect of subjective wellbeing on cardiovascular health, cholesterol or BMI. This is consistent with a prospective analysis in over 700,000 women which found no effect of happiness on later mortality, if baseline health was controlled for [55]. Previous observational associations [2,3] could be due to residual confounding, reverse causation or publication bias [56,57]. In our analysis, there was little evidence that subjective wellbeing had a causal effect on physical health outcomes. The genetic variants for subjective wellbeing are weak and confidence intervals were wide, so the null effect could be due to a lack of statistical power. Further analysis is needed when stronger instruments are available.

There was no clear evidence for a causal effect of cholesterol, coronary artery disease risk or myocardial infarction risk on subjective wellbeing, meaning that residual confounding is likely responsible for the previous observational associations. This conclusion is supported by recent evidence from a new approach called Bayesian direct multimorbidity mapping (BDMM) which found that CAD was only associated with depression because of an association with BMI [58]. However, CAD and MI are rare outcomes and the SNPs for CAD risk used in our analysis all had small effect sizes [36], resulting in limited power to detect an effect on subjective wellbeing. Future research using the continuous phenotype of blood pressure could provide additional insight into the causal pathways between cardiovascular health and subjective wellbeing.

### Limitations

In addition to the specific limitations of weak instruments and statistical power outlined above a more general limitation of this study was the use of BMI as a proxy for adiposity. BMI can vary due to reasons other than adiposity and we cannot be sure which aspect is driving the casual association. We need to understand the mechanisms clearly in order to design interventions [59]. Nevertheless, BMI is a good indicator of adiposity, is widely available and easy to collect in large samples, and other more precise measures have not been shown to differ dramatically [60].

A second possible limitation could be the influence of population structure on the genetic instrument. In large samples such as the UK Biobank, it is difficult to fully remove population structure without removing true effects [61]. Coincident structure may confound the association between BMI and subjective wellbeing, generating spurious signal. Although we cannot completely remove the possible influence of structure in our replication analysis, we are reasonably confident that the effect of BMI on satisfaction with health is not spurious because we do not see the same inflation for the negative control outcomes of domain satisfaction or happiness. Further, non-genetic instrumental variables give the same results as genetic instruments in the UK Biobank for educational attainment, a trait largely influenced by structure [62].

## Conclusion

We found no clear evidence, using MR for a causal effect of subjective wellbeing on physical health outcomes. This suggests that previously reported observational associations may have resulted from residual confounding. We found strong evidence for a causal effect of increased BMI on decreased subjective wellbeing. Replication in UK Biobank suggested that the effect of BMI on subjective wellbeing was driven by an adverse effect of higher BMI on health satisfaction. Our findings add further support to the need to reduce obesity because of the downstream consequences on mental health and wellbeing. Further work is required to understand the pathways from BMI to subjective wellbeing and to explore how the effect of BMI on mental health varies across the life-course.

## Acknowledgements

We are grateful to the participants of the UK Biobank and the individuals who contributed to the SSGAC, GLGC, CARDIoGRAMplusC4D and GIANT GWAS consortia as well as all the research staff who worked on the data collection.

## Disclaimers

The views expressed in the submitted article are those of the author and not an official position of the institution or funder.

## Contribution statement

Contributors CMAH and OSPD conceived the study. REW conducted the analysis. REW and CMAH drafted the initial manuscript. AET assisted with the two-sample MR analysis. LACM assisted with the replication analysis, created the polygenic score for BMI and provided access to the UK Biobank phenotype data. NMD provided access to the UK Biobank genetic data. RBL assisted with all sensitivity analyses. NMD, GDS, MRM, NJT, LACM and RBL advised and guided all stages of analysis. All authors assisted with interpretation, commented on drafts of the manuscript and approved the final version.

## Public Patient Involvement

None

## Ethical approval

UK Biobank has received ethics approval from the National Health Service National Research Ethics Service (ref 11/NW/0382).

## Data and code availability

Scripts are available on GitHub at: https://github.com/MRCIEU/Health-and-Wellbeing-MR

